## Supplemental Data for "A curative combination therapy for lymphomas achieves high fractional cell killing through low cross-resistance and drug additivity but not synergy"

**This document contains Figures S1 to S8 and Tables S1 and S2.**

**Tables S3 to S9 can be downloaded at:**

www.dropbox.com/sh/z39aq4rleyi4jmk/AACRxxGZekOdWgqi86szViCGa

**Following peer review these source data files will be publically shared.**

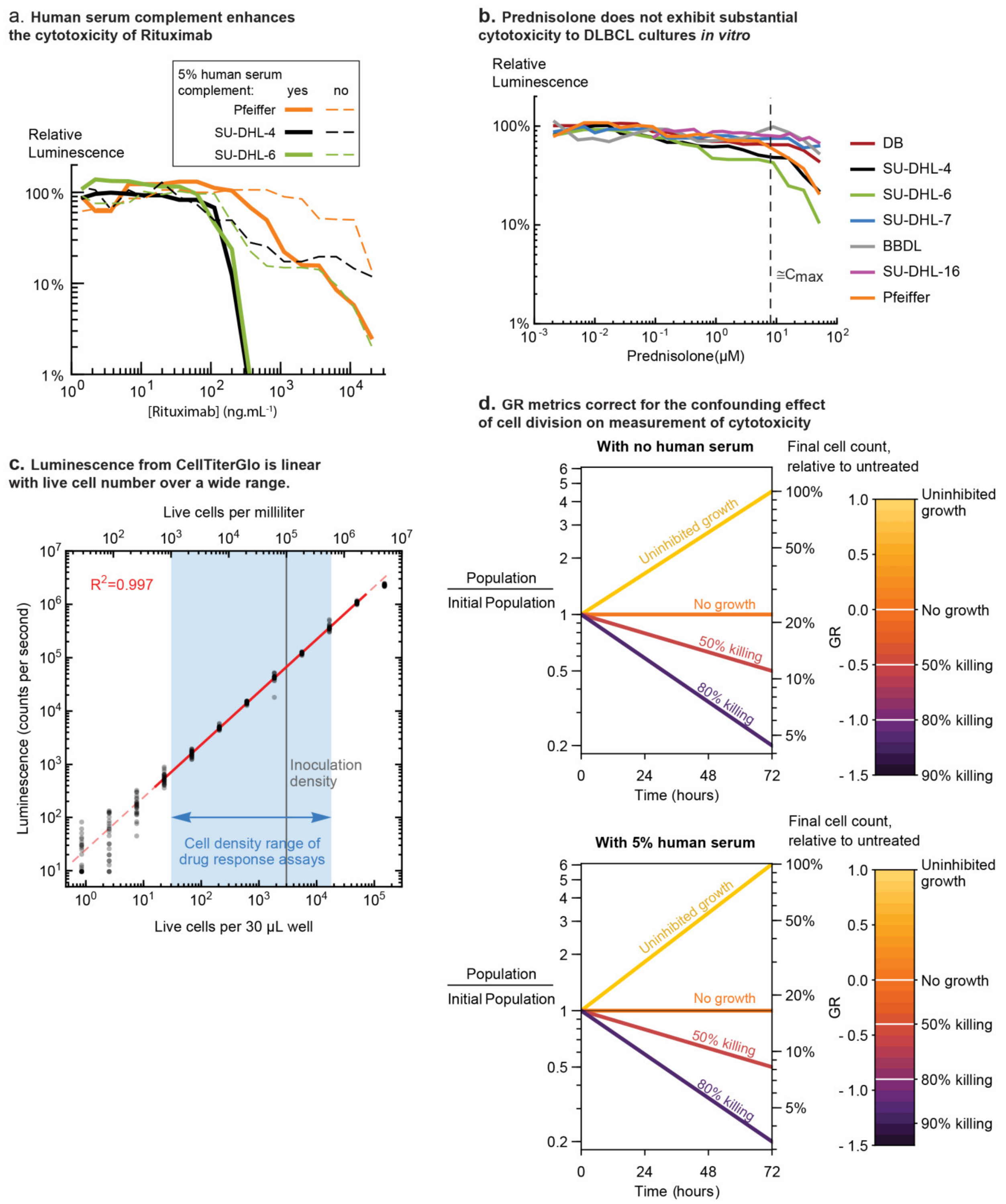


**Figure S1. Related to Figure 1. a. Human serum complement enhances the cytotoxicity of Rituximab to DLBCL cell lines.** The DLBCL cell lines Pfeiffer, SU-DHL-4, SU-DHL-6 were each seeded in 384-well plates at $10^5$ cells/mL in complete media with or without 5% pooled complement human serum, and were treated with a range of Rituximab concentrations. After 72 hours of treatment, cell viability was measured by luminescent assay of ATP (CellTiter Glo). Untreated control wells defined 100% relative luminescence. **b. Prednisolone does not exhibit substantial cytotoxicity to DLBCL cultures *in vitro*.** DLBCL cultures were seeded in 384-well plates, treated with prednisolone for 72 hours, and relative viability was assayed by CellTiter Glo. None of 7 cell lines demonstrated cytotoxicity at clinically relevant doses (dashed vertical line marks the estimated Cmax of 8 μM). Because untreated cultures divide ~2 to 3 times in 72 hours, reduction of luminescence to ~ 50% of untreated control indicates a partial reduction in proliferation rate and not net cytotoxicity (more death than growth). **c. Luminescence from CellTiter Glo is linear with live cell number over a wide range.** Pfeiffer cells were concentrated by centrifugation and resuspended in a reduced volume of media, and the density of live cells was counted using a haemocytometer and trypan blue staining. This culture was seeded in 384-well plates with a volume of 30 μL per well of complete media and cell densities ranging from $5\times10^6$ down to 50 cells per mL, prepared via serial dilution. 24 replicate wells were prepared at each density. Immediately after inoculation, the plate was subjected to a luminescent assay of ATP content by CellTiter Glo. A solid red line represents the region of highly linear relationship between live cell density and luminescence ($R^2 = 0.997$); at higher densities luminescence begins to saturate, and at lower densities significant noise is observed. A vertical gray line indicates the initial cell density used in drug response assays (Figures 1 and 2), and a blue shaded region indicates the cell density range relevant to these assays (density increasing in event of uninhibited growth, and decreasing in event of cytotoxicity). **d. Growth Rate (GR) metrics correct for the confounding effect of cell division on cytotoxicity.** Cell division during a drug treatment assay affects the relationship between cell death and cell count relative to untreated controls (Hafner et al., 2016). To correct for this effect, absolute cell counts of untreated cultures were measured before and after treatment to quantify cell proliferation (see Methods). Plots indicate the relation between relative cell number (as measured by CellTiter Glo) and fraction of cells killed. The 'no human serum' scale is relevant to pairwise interactions in CHOP (Figure 1), and '5% human serum' is relevant to pairs of R and others drugs in CHOP (Figure 1) and all measurements of high-order interactions (Figure 2). Similar relationships were measured and applied for SU-DHL-4 and SU-DHL-6.

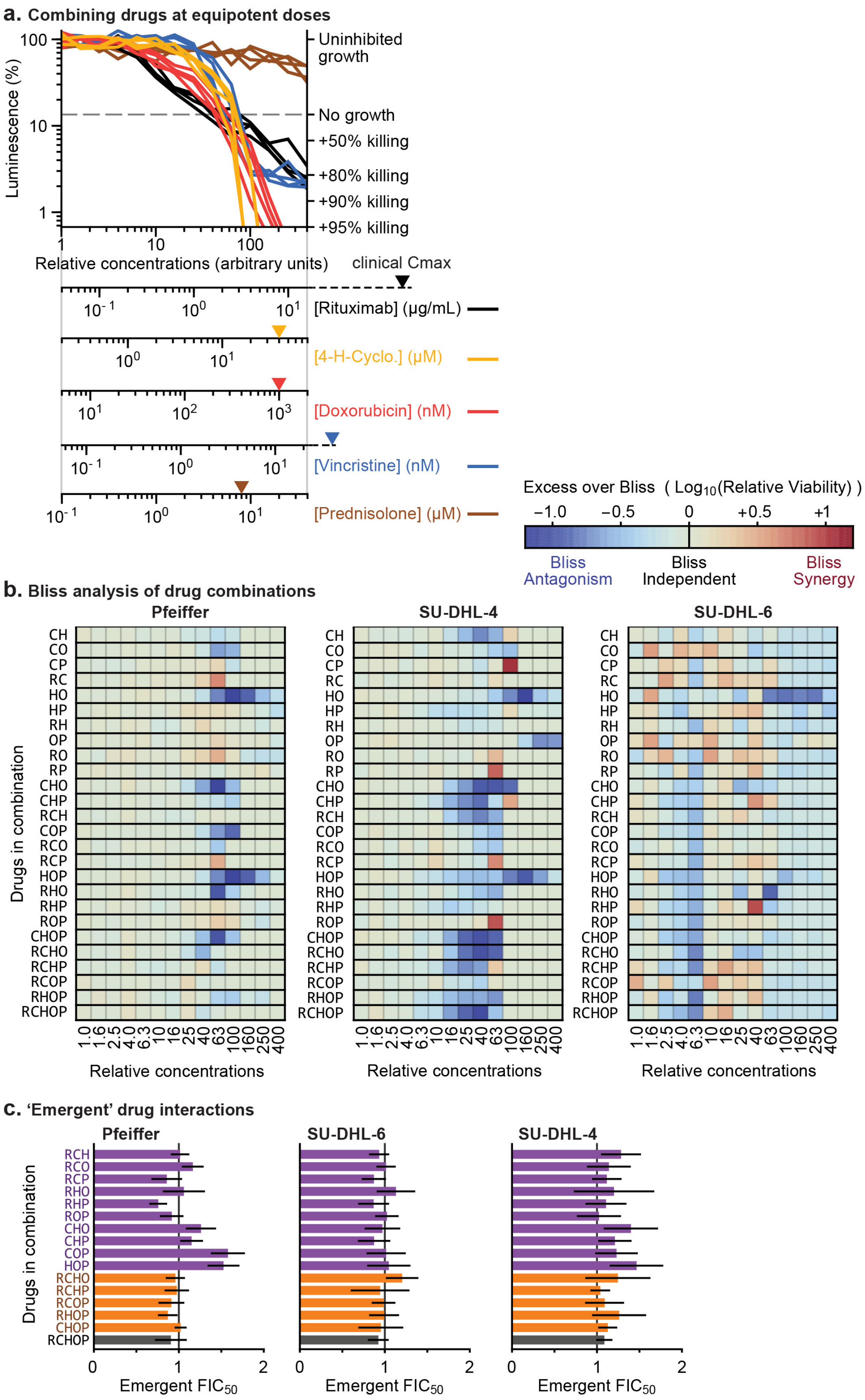
a. Combining drugs at equipotent doses
Luminescence (%)
100
10
1
Uninhibited growth
No growth
+50% killing
+80% killing
+90% killing
+95% killing
1
10
100
Relative concentrations (arbitrary units)
clinical Cmax
[Rituximab] (μg/mL)
[4-H-Cyclo.] (μM)
[Doxorubicin] (nM)
[Vincristine] (nM)
[Prednisolone] (μM)
Excess over Bliss ( Log10(Relative Viability) )
−1.0
−0.5
0
+0.5
+1
Bliss Antagonism
Bliss Independent
Bliss Synergy
b. Bliss analysis of drug combinations
Pfeiffer
SU-DHL-4
SU-DHL-6
Drugs in combination
CH
CO
CP
RC
HO
HP
RH
OP
RO
RP
CHO
CHP
RCH
COP
RCO
RCP
HOP
RHO
RHP
ROP
CHOP
RCHO
RCHP
RCOP
RHOP
RCHOP
1.0
1.6
2.5
4.0
6.3
10
16
25
40
63
100
160
250
400
Relative concentrations
c. 'Emergent' drug interactions
Pfeiffer
SU-DHL-6
SU-DHL-4
Drugs in combination
RCH
RCO
RCP
RHO
RHP
ROP
CHO
CHP
COP
HOP
RCHO
RCHP
RCOP
RHOP
CHOP
RCHOP
0
1
2
Emergent FIC50

**Figure S2. Related to Figure 2. a. Combining the drugs in RCHOP at equipotent ratio.** In order to measure high-order interactions in RCHOP (Figures 2, S2B and S2C), it was necessary to combine agents in equipotent ratio, i.e. mixing them in proportions that permit each drug to contribute similarly to cytotoxicity (except for prednisolone which does not display single-agent cytotoxicity *in vitro*). Plotted here are dose responses to each single drug as measured in the experiments of Figure 2 for Pfeiffer (n=8; including two biological duplicate cultures, here plotting four lines per drug that are each the mean of technical duplicates). For each drug a 400-fold dose range was selected (see axes below the plot) to align all 4 cytotoxic agents RCHO such that they enter the cytotoxic regime of their dose response (crossing the dashed gray line) at similar positions and that this position aligns with the clinical Cmax (peak serum concentration) of prednisolone. This alignment defines the "relative concentration" units in Figure 2C. The concentration range also aligns the clinical Cmax (colored triangles on concentration axes) of each drug, except for Rituximab which is evidently very highly potent against Pfeiffer cells. Relative luminescence (left axis) was converted to biological effect (right axis) by measuring absolute cell count of untreated cultures before and after drug treatment (Methods): in this experiment untreated cultures increased in cell count by approximately 7-fold, and therefore relative luminescence of ≈14% corresponds to no net growth or death ("no growth" on axis). A net cytotoxic effect is evidenced only by relative luminescence lower than 14% (e.g. +50% killing at ≈7% luminescence, +80% killing at ≈3% luminescence). Similar 400-fold dose windows were independently determined for SU-DHL-4 and SU-DHL-6. **b. High-order interactions of RCHOP analyzed according to the Bliss Independence model in 3 DLBCL cell lines using data from the experiment displayed in Figure 2.** The Bliss Independence model defines toxins to be 'non-interacting' when they confer statistically independent probabilities of cell death, thus, surviving fractions of cells are multiplied (e.g. 90% inhibition by drug A + 90% inhibition by drug B should produce 99% inhibition in combination). The 'Excess over Bliss' measures the observed deviation from the Bliss Independence model: in these heat maps blue indicates antagonism (less killing), and red indicates synergy (more killing). See legend above the 'SU-DHL-6' graph. The combination of all 5 drugs in RCHOP shows neither synergy nor antagonism, being consistent with Bliss independence (bottom row of panels). **c. Emergent high-order interactions of RCHOP analyzed using fractional inhibitory concentrations at 50% killing threshold ($FIC_{50}$) in 3 DLBCL cell lines.** Emergent FIC quantifies any deviation in the potency of 3, 4, or 5-way combinations from the assumption of additivity between known lower-order interactions (Cokol et al., 2017). FIC is the sum of each drug's concentration in that mix as a fraction of their single-agent dose producing the same effect: $FIC_{50} = \sum \frac{IC50_{\text{drug in combination}}}{IC50_{\text{drug alone}}}$; and Emergent FIC $= FIC_{N\text{ drugs}} \,/\, \overline{FIC_{N-1\text{ drugs}}}$ (e.g. for three drugs ABC: $FIC_{ABC}$ / average( $FIC_{AB}$, $FIC_{AC}$, $FIC_{BC}$ ). Error bars are 95% confidence intervals (n=8 per dose point for single-drug dose responses (denominator), n=4 per dose point for multi-drug dose responses (numerator)).

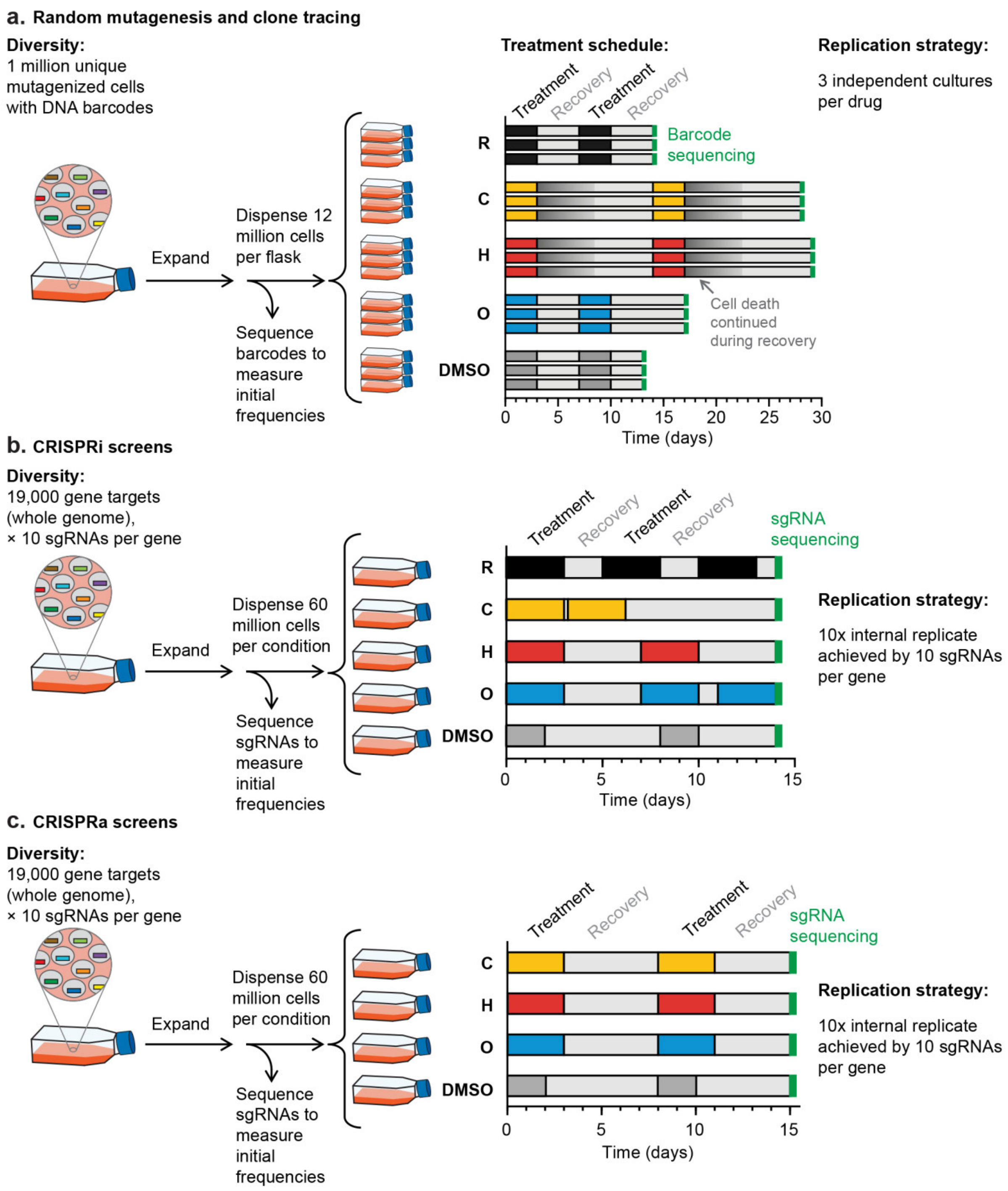


**Figure S3. Related to Figure 3. Dosing schedule and replication strategy for all 3 approaches taken to isolate resistant cells to single cytotoxic drugs in the RCHOP combination. a.** Experimental design for measuring drug resistance in DNA-barcoded Pfeiffer clones. 1 million unique mutagenized Pfeiffer cells individually tagged with DNA barcodes were expanded. Replicate flasks of cells (3 per drug) were treated with 2 pulses of drug according to the dosing regimen depicted. Recovery was slowest after treatment by C or H, which induced sustained cell death for days after medium was exchanged and drug removed. The abundance of each clone was determined in the pre-treatment population and in each drug-treated replicate sample by high-throughput sequencing. **b.** Experimental design for measuring drug resistance in a whole-genome collection of knockdown mutants (CRISPRi). Pfeiffer CRISPRi cells transduced with a genome-wide library of sgRNAs were treated with 2-3 72 h pulses of drug according to the dosing regimen depicted. The treatment schedule was adjusted in order to achieve 7-9 fewer population doublings than the DMSO control. The CRISPRi library contains 10 sgRNAs per gene which serve as internal replicates for each data point. **c.** Experimental design for measuring drug resistance in a whole-genome collection of overexpression mutants (CRISPRa). K562 CRISPRa cells transduced with a genome-wide library of sgRNAs were treated with two 72 h pulses of drug according to the dosing regimen depicted. The treatment schedule was adjusted in order to achieve 9-10 fewer population doublings than the DMSO control. The CRISPRa library contains 10 sgRNAs per gene which serve as internal replicates for each data point.

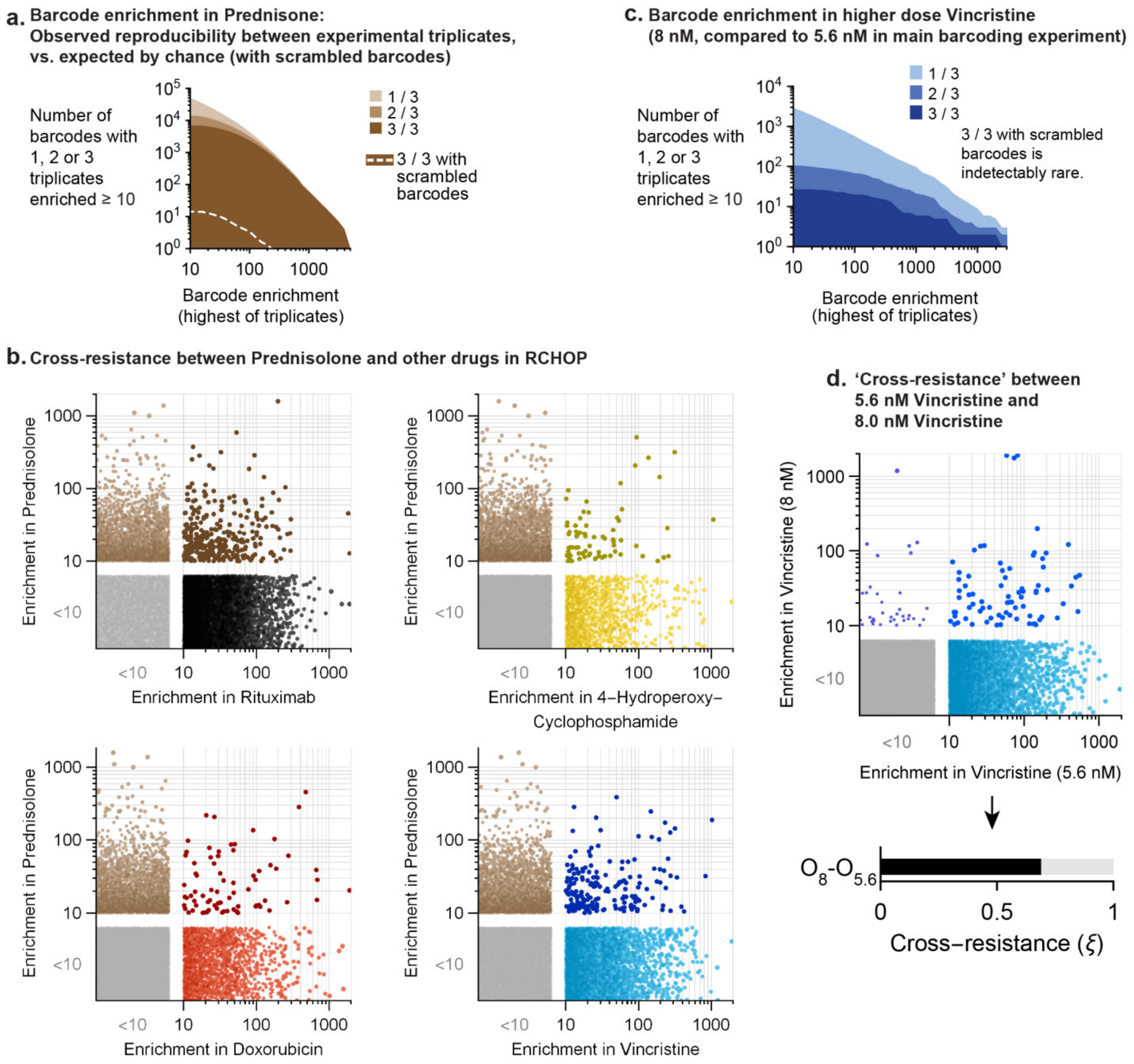


**Figure S4. Related to Figure 4. Prednisolone-resistant clones show low cross-resistance to other drugs in RCHOP, and repeats of vincristine show high cross-resistance. a**. Enriched barcodes in the cultures treated with prednisolone show a high degree of reproducibility between 3 replicate drug selections. As prednisolone did not show single-agent cytotoxicity but only partially inhibited growth (Figure S1B), barcoded Pfeiffer cultures were exposed to 20 days of continuous treatment with 20µM prednisolone (see Methods). The number of barcodes reproducibly enriched by prednisolone treatment was far in excess (> 300-fold) of the number of barcodes expected by random chance (error model based on scrambled barcode identities; Methods). **b**. Scatter plots displaying enrichment scores in prednisolone and other drugs in RCHOP. Enrichment scores of individual barcodes were determined by the geometric mean enrichment of triplicates, and were deemed significant at a score above 10 (based on false discovery rate of double-resistant clones). **c**. A set of biological triplicate barcoding experiments were conducted at a higher dose of vincristine (8 nM) than was used in the main barcoding experiments (5.6 nM vincristine). The higher dose imposed far stronger selection: the population required ≈2 weeks to recover following each of two 72 h treatment cycles, and fewer barcodes exhibited significant enrichment. **d**. Analysis of 'cross-resistance' between the two doses of vincristine. The majority of barcodes (≈70%) enriched in 8 nM vincristine were also enriched in 5.6 nM vincristine (the opposite cannot be true because fewer barcodes in total were enriched at 8 nM). This corresponds to a cross-resistance parameter of ξ=0.69, a value that indicates close to maximally overlapping resistance.

a.

log2 fold change in expression CRISPRi controls

| qPCR primers: | sgRNA targeting: ST3GAL4 | SEL1L | DPH1 | Non-targeting |
|---|---|---|---|---|
| ST3GAL4 | -3.9 ± 0.5 | 0.6 ± 0.3 | 0.0 ± 0.4 | 0.0 ± 0.1 |
| SEL1L | 0.3 ± 0.9 | -2.9 ± 0.3 | -0.6 ± 0.8 | 0.0 ± 0.3 |
| DPH1 | -0.5 ± 0.5 | -1.2 ± 0.1 | -5.4 ± 0.8 | 0.0 ± 0.3 |

log2 fold change in expression CRISPRa controls

| qPCR primers: | sgRNA targeting: CDKN1C | SLC4A1 | POU5F1 | Non-targeting |
|---|---|---|---|---|
| CDKN1C | 6.7 ± 0.8 | 0.3 ± 0.4 | 0.1 ± 0.7 | 0.0 ± 0.4 |
| SLC4A1 | 4.2 ± 1.2 | 8.1 ± 0.4 | 0.4 ± 0.9 | 0.0 ± 2.0 |
| POU5F1 | 0.5 ± 0.8 | 1.1 ± 1.5 | 11.5 ± 0.5 | 0.0 ± 0.2 |

b.

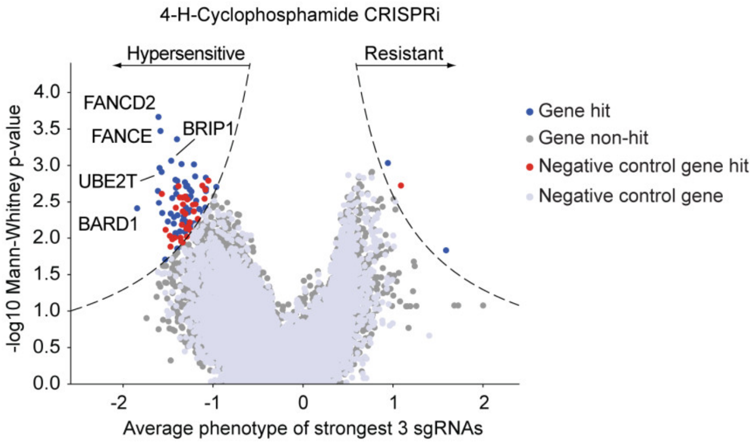


c.

Hypersensitive genes GO term enrichment analysis:

| Description | P-value | FDR q-value |
|---|---|---|
| interstrand cross-link repair | 1.32E-14 | 2.02E-10 |
| DNA repair | 6.05E-09 | 4.62E-05 |
| meiotic cell cycle process | 1.54E-07 | 7.82E-04 |
| DNA metabolic process | 8.50E-07 | 2.60E-03 |
| cellular response to DNA damage stimulus | 7.56E-07 | 2.89E-03 |
| homophilic cell adhesion via plasma membrane adhesion molecules | 2.33E-05 | 3.96E-02 |
| protein K6-linked ubiquitination | 2.12E-05 | 4.06E-02 |
| cell-cell adhesion via plasma-membrane adhesion molecules | 1.99E-05 | 4.34E-02 |
| double-strand break repair | 1.93E-05 | 4.92E-02 |
| cell cycle checkpoint | 3.42E-05 | 5.22E-02 |
| biological adhesion | 6.15E-05 | 8.55E-02 |
| regulation of cell cycle process | 7.42E-05 | 9.45E-02 |

Genes associated with the interstrand cross-link repair GO term found in screen:
FANCE - fanconi anemia, complementation group e
FANCD2 - fanconi anemia, complementation group d2
ERCC4 - excision repair cross-complementing rodent repair deficiency, complementation group 4
UBE2T - ubiquitin-conjugating enzyme e2t (putative)
EME1 - essential meiotic structure-specific endonuclease 1
RAD51D - rad51 paralog d
FANCA - fanconi anemia, complementation group a
FANCI - fanconi anemia, complementation group i
FANCL - fanconi anemia, complementation group l
ATRIP - atr interacting protein

**Figure S5. Related to Figure 5. CRISPRi/a cell lines strongly alter gene expression of targeted genes and additional cyclophosphamide CRISPRi screen identifies hypersensitive hits in the DNA interstrand crosslink pathway. a.** CRISPRi targeting of 3 control genes results in knockdown in Pfeiffer cells (left) and CRISPRa targeting of 3 control genes results in overexpression in K562 cells (left). On-target effect is strong (green shaded boxes) whereas off-target and/or indirect effects are low. Cell lines expressing either the CRISPRi or CRISPRa system were transduced with sgRNAs targeting control genes. The expression of control genes was measured in those cell lines by RT-qPCR using gene-specific primers and was compared to a non-targeting control sgRNA (see Methods). **b.** Volcano plot of gene phenotype and enrichment p-value for the CRISPRi screen of 4-H-Cyclophosphamide performed at a lower concentration (as compared to Figure 5). The position of each gene was determined by the average phenotype of the 3 most active sgRNAs (out of a library of 5sgRNAs per gene) and $-\log_{10}$ of the p-value determined using the Mann-Whitney test of all sgRNAs targeting that gene compared to non-targeting controls. For genes with multiple transcription start sites (TSSs) targeted, sgRNAs were grouped by TSS and the TSS with the lowest p-value was used for display and downstream analysis. Negative control genes were generated by randomly grouping sets of 5 non-targeting controls which were subsequently analyzed as true genes. Dashed lines represents a cutoff $\geq 7$ as calculated from: (gene phenotype z-score)$\times$($-\log_{10}$(p-value)) and where the z-score is calculated from the standard deviation of the set of negative control genes. **c.** Table containing the top hits from a gene ontology analysis performed on the CRISPRi hypersensitive hits. A hypersensitivity score was calculated for each gene by multiplying the two metrics determined in the screen (score = $-\log_{10}$(p-value)$\times$gene phenotype). Gene ontology analysis was performed on the list of genes ranked by hypersensitivity score using GOrilla (Eden et al., 2009). Reported p-values are the enrichment p-values computed according to the GOrilla algorithm and the 'FDR q-values' represent the correction of the p-values for multiple testing hypothesis. A list of all genes belonging to the GO term 'interstrand cross-link repair' and which were found as hypersensitive in the CRISPRi cyclophosphamide screen is reported below the table.

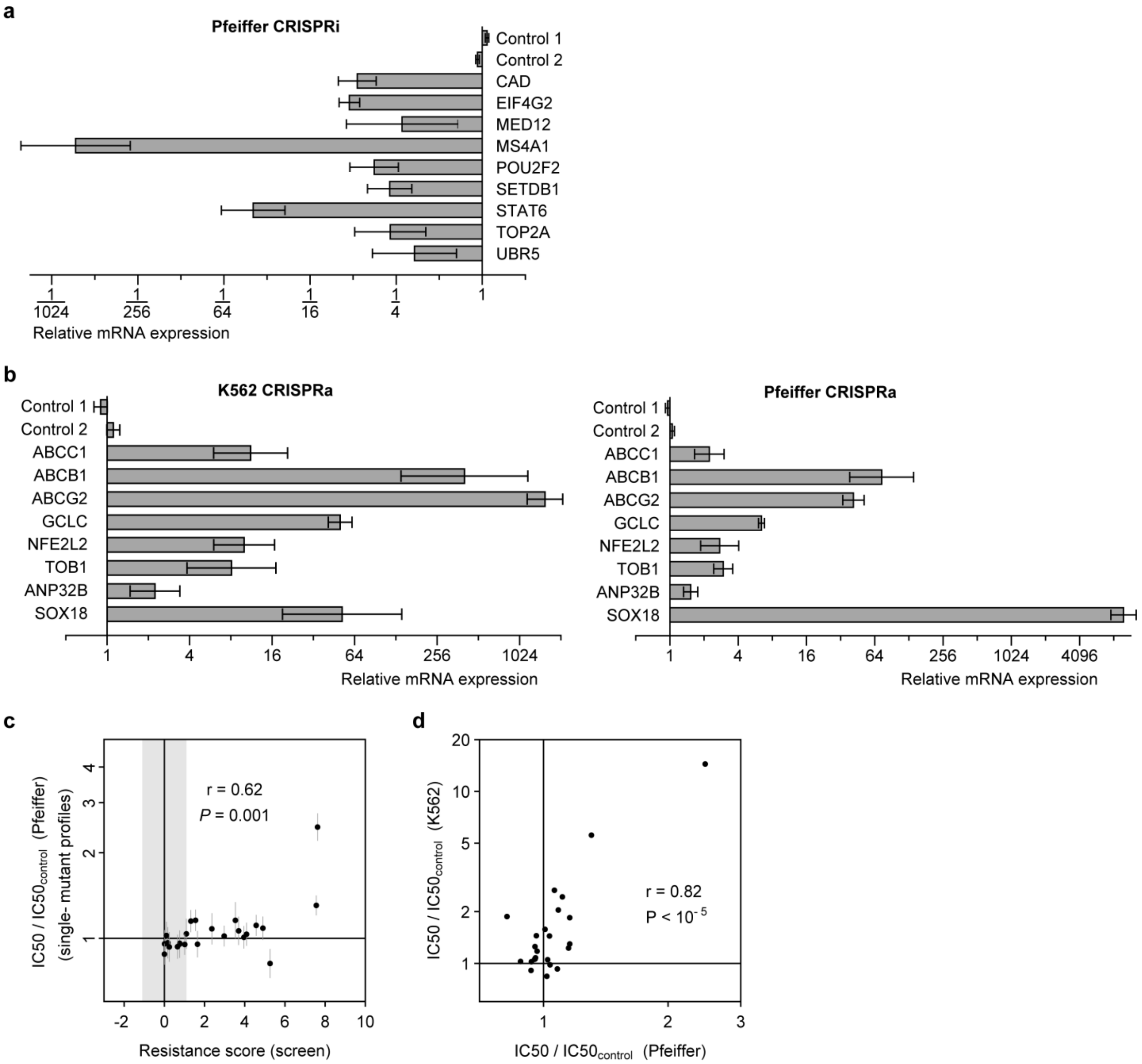


**Figure S6. Related to Figure 6. (a-b) Quantification of changes in gene expression in validation CRISPRi and CRISPRa cell lines and (c-d) Gene overexpression by CRISPRa produces changes in drug resistance that are correlated across K562 and Pfeiffer DLBCL cells, but weaker in Pfeiffer. a.** Measurement of changes in mRNA levels by RT-qPCR in Pfeiffer CRISPRi cells transduced with individual sgRNAs for each of 9 knockdown screen hits. Controls 1 and 2 represent cell lines expressing non-targeting sgRNAs and the average of the two cell lines was used for normalization. Error bars are 95% confidence intervals (n=3 technical repeats). **b**. same as **a.** except for K562 CRISPRa cell lines (left) and Pfeiffer CRISPRa cell lines (right). **c.** Drug dose response measurements in Pfeiffer CRISPRa cells transduced with individual sgRNAs, for each of 8 overexpression screen hits, show changes in IC50 that are correlated with resistance phenotypes measured in the genome-wide CRISPRa screen in K562, but weak in their magnitude of effect in Pfeiffer. Error bars are 95% confidence intervals in IC50 (determined from curve fit; n=3 biological repeats). Gray region: threshold in resistance score that was used to identify screen hits. **d**. For the same screen hits in panel c, gene overexpression by CRISPRa produces changes in IC50 in Pfeiffer DLBCL cells that are correlated but weaker than changes in IC50 measured in K562, where CRISPRa produces stronger induction of gene expression.

a.

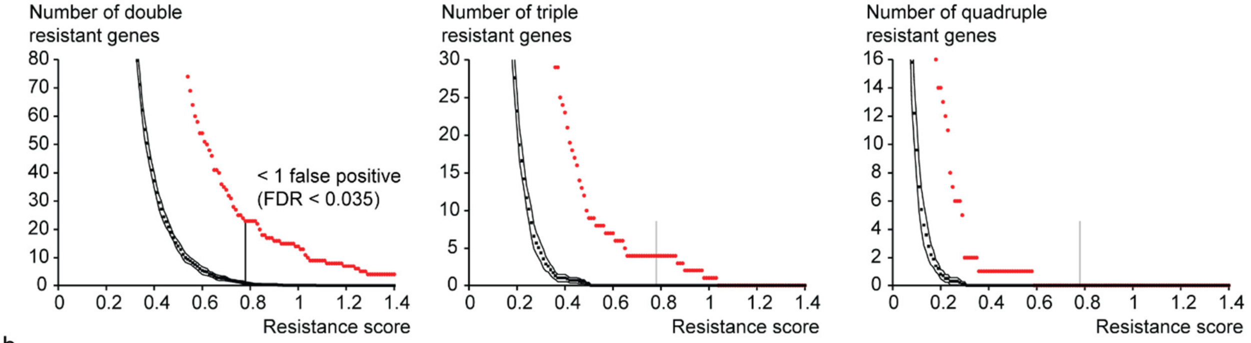


b.

| | R | C | H |
|---|---|---|---|
| C | EIF4G2, DBR1, SETDB1, BAHD1, MED12 | | |
| H | EIF3G, NAPA, EIF4G2, SMN2, UBR5, MED12, MED16, CAD, SMARCE1, FNBP4 | MIOS, UBA52, RPS28, TOP2A, ATP5J2, EIF4G2, POU2F2, ATP5J2-PTCD1, MED12 | |
| O | SETDB1, UBR5 | SETDB1, MAU2 | RBM12, CCNC, UBR5 |

EIF4G2: Eukaryotic translation initiation factor 4 gamma 2
SETDB1: SET domain bifurcated 1 (histone-lysine N-methyltransferase)
MED12: Mediator complex subunit 12
UBR5: Ubiquitin protein ligase E3 component n-recognin 5

c.

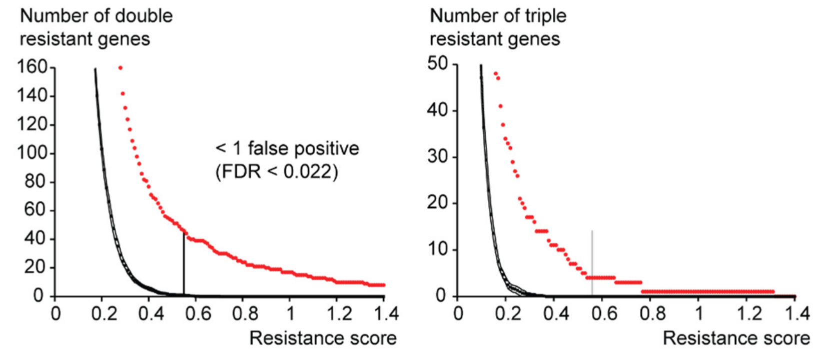


d.

| | O | C |
|---|---|---|
| C | BAG6, RUNX1T1, FGFR1, CHD7, FLI1, PGC, CUL1, COL4A3BP, GLS, CBFA2T2, ZNF592, POLK, RUNX2, NFE2L2, HIAT1, MED24, DSTYK, SOX18, TOB1, NR4A1, ANP32B, GAREM, GCLC, JUN, LDLR, ZNF608, RCOR2, SIX4 | |
| H | ACTR1A, TMEM150A, EEPD1, FAM89B, FOPNL, RAB7A, TOB1, GATA2, ANP32B, NR5A2, GCLC, OSGIN1, ABCB1, ALAS2, ABCC1, NFE2L2, ATP10A, | SIK2, PEG10, SESN2, FOXO3, GCLC, TOB1, MOV10, NFE2L2, ANP32B |

NFE2L2: Nuclear factor, erythroid 2 like 2
GCLC: glutamate-cysteine ligase catalytic subunit
TOB1: transducer of ERBB2, 1
ANP32B: Acidic nuclear phosphoprotein 32 family member B

**Figure S7. Related to Figure 7. Determination of a cutoff threshold for cross-resistance analysis and identity of all cross-resistant genes for CRISPRi and CRISPRa screens. a.** Plots of the total number of double-, triple- and quadruple-resistant genes as a function of the resistance score determined in the CRISPRi screens. Whole genome sets of negative control genes were simulated by randomly grouping sets of non-targeting sgRNAs. The resistance score for each negative control gene was then calculated for each drug CRISPRi screen. Black dots represent the average number of negative control genes (over a total of 10 simulated sets) that score as cross-resistant as a function of the resistance score. Black lines represent 90% confidence interval lines. Red dots represent the number of true cross-resistant genes. The threshold for cross-resistance analysis was set at a score that yielded less than an average of one double-resistant negative control gene. The false positive rate (FDR) was calculated by dividing the average number of double-resistant negative control genes by the number of true double-resistant genes. **b.** Cross-resistant genes identified in the CRISPRi screens for RCHO. Genes that are triple-resistant are highlighted in colored font. **c.** Plots of the total number of double- and triple-resistant genes as a function of the resistance score determined in the CRISPRa screens. Data were analyzed and displayed as for a. **d.** Cross-resistant genes identified in the CRISPRa screens for CHO. Genes that are triple-resistant are highlighted in red.

a.

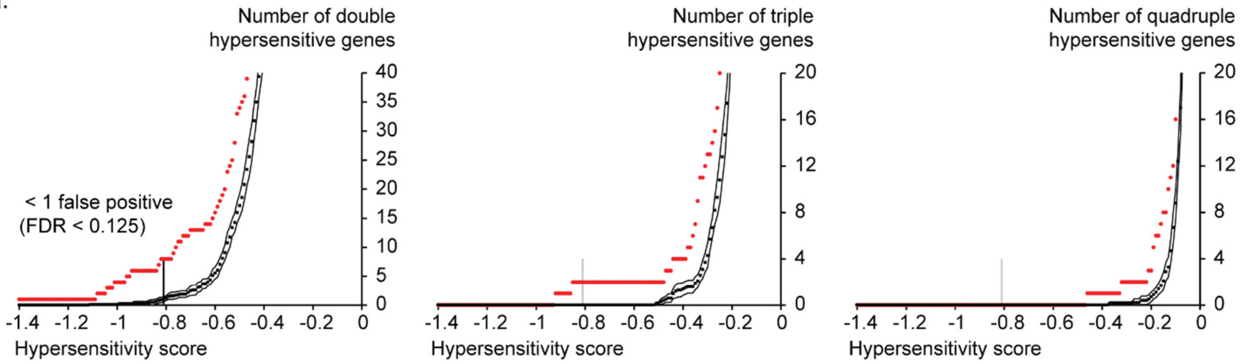


b.

| | R | C | H |
|---|---|---|---|
| C | - | | |
| H | DDX6, PPP2R5C, NPRL3 | - | |
| O | DDX6, PPP2R5C | BRCA2 | SLC38A10, DDX3X, GCLC, PPP2R5C, DDX6, GIGYF2, |

DDX6: DEAD-box helicase 6
PPP2R5C: Protein phosphatase 2 regulatory subunit B' gamma

c.

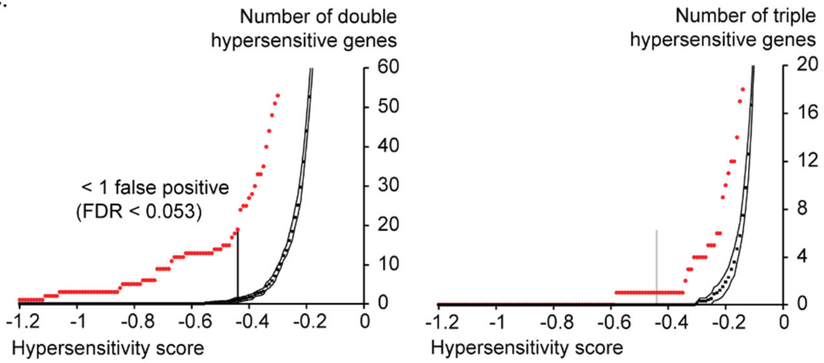


d.

| | O | C |
|---|---|---|
| C | BCL9L, SSBP2, LMO2, ZNF740, CTDSPL2, TP53BP1, TRPV3 | |
| H | ZMIZ1, BCL9L, MTDH, FRAT2, SLC1A5, HNRNPU, CD44, SLCO4A1, KLK11 | ETS2, BCL9L, ETS1, FAM60A, CKS2 |

BCL9L: B cell CLL/lymphoma 9 like

**Figure S8. Related to Figure 7. Determination of a cutoff threshold for cross-hypersensitivity analysis and identity of all cross-hypersensitive genes for CRISPRi and CRISPRa screens. a.** Plots of the total number of double-, triple- and quadruple-hypersensitive genes as a function of the hypersensitivity score determined in the CRISPRi screens. Whole genome sets of negative control genes were simulated by randomly grouping sets of non-targeting sgRNAs. The hypersensitivity score for each negative control gene was then calculated for each drug CRISPRi screen. Black dots represent the average number of negative control genes (over a total of 10 simulated sets) that score as cross-hypersensitive as a function of the hypersensitivity score. Black lines represent 90% confidence interval lines. Red dots represent the number of true cross-hypersensitive genes. The threshold for cross-hypersensitivity analysis was set at a score that yielded less than an average of one double-hypersensitive negative control gene. The false positive rate (FDR) was calculated by dividing the average number of double-hypersensitive negative control genes by the number of true double-hypersensitive genes. **b.** Cross-hypersensitive genes identified in the CRISPRi screen for RCHO. Genes that are triple-hypersensitive are highlighted in red font. **c.** Plots of the total number of double- and triple-hypersensitive genes as a function of the hypersensitivity score determined in the

CRISPRa screens. Data were analyzed and displayed as for a. **d.** Cross-hypersensitive genes identified in the CRISPRa screen for CHO. Genes that are triple-hypersensitive are highlighted in red.

| Oligo Name | Sequence | Description |
| --- | --- | --- |
| CC_LSP_001 | TTGGGGCGCGGGTCCGGCCTGGGAGGTTTAAGAGC | ST3GAL4 sgRNA construction |
| CC_LSP_002 | TTAGCTCTTAAACCTCCCAGGCCGGACCCGCGCCCCAACAAG | ST3GAL4 sgRNA construction |
| CC_LSP_003 | TTGGCAGGAAGAGCAGCGGCGAGGGTTTAAGAGC | SEL1L sgRNA construction |
| CC_LSP_004 | TTAGCTCTTAAACCCTCGCCGCTGCTCTTCCTGCCAACAAG | SEL1L sgRNA construction |
| CC_LSP_005 | TTGGTATCCGGGGCAGCGGAGCAGTTTAAGAGC | DPH1 sgRNA construction |
| CC_LSP_006 | TTAGCTCTTAAACTGCTCCGCTGCCCCGGATACCAACAAG | DPH1 sgRNA construction |
| CC_LSP_079 | TTGGCTGCATGGGGCGCGAATCAGTTTAAGAGC | Non targeting sgRNA construction |
| CC_LSP_080 | TTAGCTCTTAAACTGATTCGCGCCCCATGCAGCCAACAAG | Non targeting sgRNA construction |
| CC_LSP_007 | TTGGCCGGGGCGCGCGGCTGATTGGGTTTAAGAGC | CDKN1C sgRNA construction |
| CC_LSP_008 | TTAGCTCTTAAACCCAATCAGCCGCGCGCCCCGGCCAACAAG | CDKN1C sgRNA construction |
| CC_LSP_009 | TTGGTCAGGAGAACCATGGGGACCGTTTAAGAGC | SLC4A1 sgRNA construction |
| CC_LSP_010 | TTAGCTCTTAAACGGTCCCCATGGTTCTCCTGACCAACAAG | SLC4A1 sgRNA construction |
| CC_LSP_011 | TTGGGATGTTTGCCTAATGGTGGGTTTAAGAGC | POU5F1 sgRNA construction |
| CC_LSP_012 | TTAGCTCTTAAACCCACCATTAGGCAAACATCCCAACAAG | POU5F1 sgRNA construction |
| CC_LSP_113 | TTGGTCATCAAGGAGCATTCCGTGTTTAAGAGC | Non targeting sgRNA construction |
| CC_LSP_114 | TTAGCTCTTAAACACGGAATGCTCCTTGATGACCAACAAG | Non targeting sgRNA construction |
| CC_LSP_013 | GATCACGCTCAAGTCCATGG | ST3GAL4 qPCR fwd |
| CC_LSP_014 | CTTGCCCAGGTCAGAAGGA | ST3GAL4 qPCR rev |
| CC_LSP_015 | GAGGGGGAAAGTGTCACAGA | SEL1L qPCR fwd |
| CC_LSP_016 | GGTCAAAGCTGGTTTCCGTA | SEL1L qPCR rev |
| CC_LSP_133 | TGGAGGCCGTTGTGTATCTT | DPH1 qPCR fwd |
| CC_LSP_134 | GACATTGGGGTTGGCAATCA | DPH1 qPCR rev |
| GAPDH78for | GGACTCATGACCACAGTCCA | GAPDH qPCR fwd |
| GAPDH78rev | GATGTTCTGGAGAGCCCCG | GAPDH qPCR rev |
| CC_LSP_143 | TGATCTCCGATTTCTTCGCCA | CDKN1C qPCR fwd |
| CC_LSP_144 | CTAAATTGGCTCACCGCAGC | CDKN1C qPCR rev |
| CC_LSP_135 | TTGCGTTCCGAGTTTCCCAT | SLC4A1 qPCR fwd |
| CC_LSP_136 | ATCTTGCGCTCAGGTTTGCT | SLC4A1 qPCR rev |
| CC_LSP_137 | AAACCCACACTGCAGCAGAT | POU5F1 qPCR fwd |
| CC_LSP_138 | TAGTCGCTGCTTGATCGCTT | POU5F1 qPCR rev |
| CC_LSP_025 | AATGATACGGCGACCACCGAACACTCTTTCCCTACACGACGC<br>TCTTCCGATCTCCTTGGAGAACCACCTTGTTG | Crispr screen fwd |
| CC_LSP_026 | AATGATACGGCGACCACCGAACACTCTTTCCCTACACGACGC<br>TCTTCCGATCTACCTTGGAGAACCACCTTGTTG | Crispr screen fwd |
| CC_LSP_027 | AATGATACGGCGACCACCGAACACTCTTTCCCTACACGACGC<br>TCTTCCGATCTAGCCTTGGAGAACCACCTTGTTG | Crispr screen fwd |
| CC_LSP_028 | AATGATACGGCGACCACCGAACACTCTTTCCCTACACGACGC<br>TCTTCCGATCTGAGCCTTGGAGAACCACCTTGTTG | Crispr screen fwd |
| CC_LSP_029 | AATGATACGGCGACCACCGAACACTCTTTCCCTACACGACGC<br>TCTTCCGATCTTTAGCCTTGGAGAACCACCTTGTTG | Crispr screen fwd |
| CC_LSP_030 | AATGATACGGCGACCACCGAACACTCTTTCCCTACACGACGC<br>TCTTCCGATCTGTAGACCTTGGAGAACCACCTTGTTG | Crispr screen fwd |

| | | |
|---|---|---|
| CC_LSP_031 | AATGATACGGCGACCACCGAACACTCTTTCCCTACACGACGCTCTTCCGATCTCAGAAACCTTGGAGAACCACCTTGTTG | Crispr screen fwd |
| CC_LSP_032 | AATGATACGGCGACCACCGAACACTCTTTCCCTACACGACGCTCTTCCGATCTTGTAGAACCTTGGAGAACCACCTTGTTG | Crispr screen fwd |
| CC_LSP_033 | CAAGCAGAAGACGGCATACGAGATTACAAGGTGACTGGAGTTCAGACGTGTGCTCTTCCGATCCGACTCGGTGCCACTTTTTC | Crispr screen rev index 1 |
| CC_LSP_034 | CAAGCAGAAGACGGCATACGAGATATTGGCGTGACTGGAGTTCAGACGTGTGCTCTTCCGATCCGACTCGGTGCCACTTTTTC | Crispr screen rev index 2 |
| CC_LSP_035 | CAAGCAGAAGACGGCATACGAGATGGAACTGTGACTGGAGTTCAGACGTGTGCTCTTCCGATCCGACTCGGTGCCACTTTTTC | Crispr screen rev index 3 |
| CC_LSP_036 | CAAGCAGAAGACGGCATACGAGATAAGCTAGTGACTGGAGTTCAGACGTGTGCTCTTCCGATCCGACTCGGTGCCACTTTTTC | Crispr screen rev index 4 |
| CC_LSP_037 | CAAGCAGAAGACGGCATACGAGATGCCTAAGTGACTGGAGTTCAGACGTGTGCTCTTCCGATCCGACTCGGTGCCACTTTTTC | Crispr screen rev index 5 |
| CC_LSP_038 | CAAGCAGAAGACGGCATACGAGATCGTGATGTGACTGGAGTTCAGACGTGTGCTCTTCCGATCCGACTCGGTGCCACTTTTTC | Crispr screen rev index 6 |
| CC_LSP_039 | CAAGCAGAAGACGGCATACGAGATCCACTCGTGACTGGAGTTCAGACGTGTGCTCTTCCGATCCGACTCGGTGCCACTTTTTC | Crispr screen rev index 7 |
| CC_LSP_040 | CAAGCAGAAGACGGCATACGAGATTTGACTGTGACTGGAGTTCAGACGTGTGCTCTTCCGATCCGACTCGGTGCCACTTTTTC | Crispr screen rev index 8 |

**Table S1. List of oligonucleotides used in this study.**

| Gene | Gamma growth scores (i.e. DMSO vs T=0) | | | |
|---|---|---|---|---|
| | **Transcript** | **Mann-Whitney p-value** | **Average phenotype of strongest 5** | **# of sgRNAs passing read count filter** |
| CAD | P1P2 | 5.89E-08 | -0.31232 | 10 |
| EIF4G2 | P1 | 1.40E-05 | -0.28846 | 10 |
| SETDB1 | P1P2 | 1.79E-06 | -0.24030 | 10 |
| MED12 | P1P2 | 8.93E-07 | -0.25856 | 10 |
| UBR5 | P1P2 | 2.08E-07 | -0.22833 | 10 |

**Table S2. Gamma growth scores for triple-resistant genes identified in CRISPRi screens.** This data was extracted from Table S8.
